# A recombinant norrin-derived growth factor retains the complete multifunctionality of human norrin

**DOI:** 10.1101/2025.10.15.682582

**Authors:** Wendy Dailey, Kenneth Mitton, Kimberly Drenser

## Abstract

Current therapies for ischemic retinopathies address the disrupted vasculature however they do not regrow patent vessels or directly benefit the neural retina. VEGF blocking therapies temporarily reduce the retinal edema but do not counter end-stage fibrosis that can lead to vision threatening consequences such as retinal detachment. Norrin is a multifunctional growth factor that is essential for neurosensory development in the eye and ear. We developed a norrin mimetic, Noregen, to overcome the deficits of currently available therapies for retinal ischemic retinopathies. Here we show that Noregen is as good or better than norrin at shoring up the retinal vascular barrier, growing patent vessels, activating neurogenic pathways and inhibiting fibrotic pathways.

## Introduction

Norrin protein plays a multifunctional role in the normal development and maintenance of central nervous system (CNS) tissues.^1,2^ Uniquely, in addition to functioning as a cysteine-knot growth factor it also serves as a Wnt ligand for multiple Wnt receptors.^3^ Norrin modulates the growth of non-fenestrated vessels, establishes and maintains the blood-retina(brain)-barrier, provides neuroprotection, stimulates neurogenesis and prevents proliferative fibrosis in the retina.^4–7^ An interplay of various receptors, receptor complexes, and protein agonism and antagonism results in these coordinated pathway activities. Alterations affecting these pathways present clinically as a spectrum of phenotypes and characteristics such as incomplete vascularization of the retina, capillary drop-out, breakdown of the blood-retina-barrier, abnormal angiogenesis and peri-retinal fibrosis. Various antibody and fusion protein strategies targeting elements of this biology are currently under investigation, but harnessing the multifunctionality of norrin in this way is problematic since the diverse scope of norrin’s receptor interactions and growth factor activities exceeds the design parameters of such agents. Therefore, we set out to mimic human norrin by creating a recombinant norrin-derived growth factor, Noregen, that could preserve all of norrin’s functionality and be scaled for GMP manufacturing (Fig. 1). Noregen was engineered and expressed in *E. coli* cells, and a proprietary manufacturing protocol for proper protein folding and dimerization was developed and validated in collaboration with Wacker Biotech GmbH (Halle, Germany). To validate that Noregen elicited proper activity and maintained the multifunctionality of human norrin, we evaluated key aspects of known norrin behavior: 1) endothelial barrier genesis; 2) retinal vascular regeneration; 3) neurogenic pathway activation, and 4) anti-fibrosis. Here we show that Noregen mimics the multiple activities of human norrin and is equivalent or superior in these functions.

**Figure 1.**
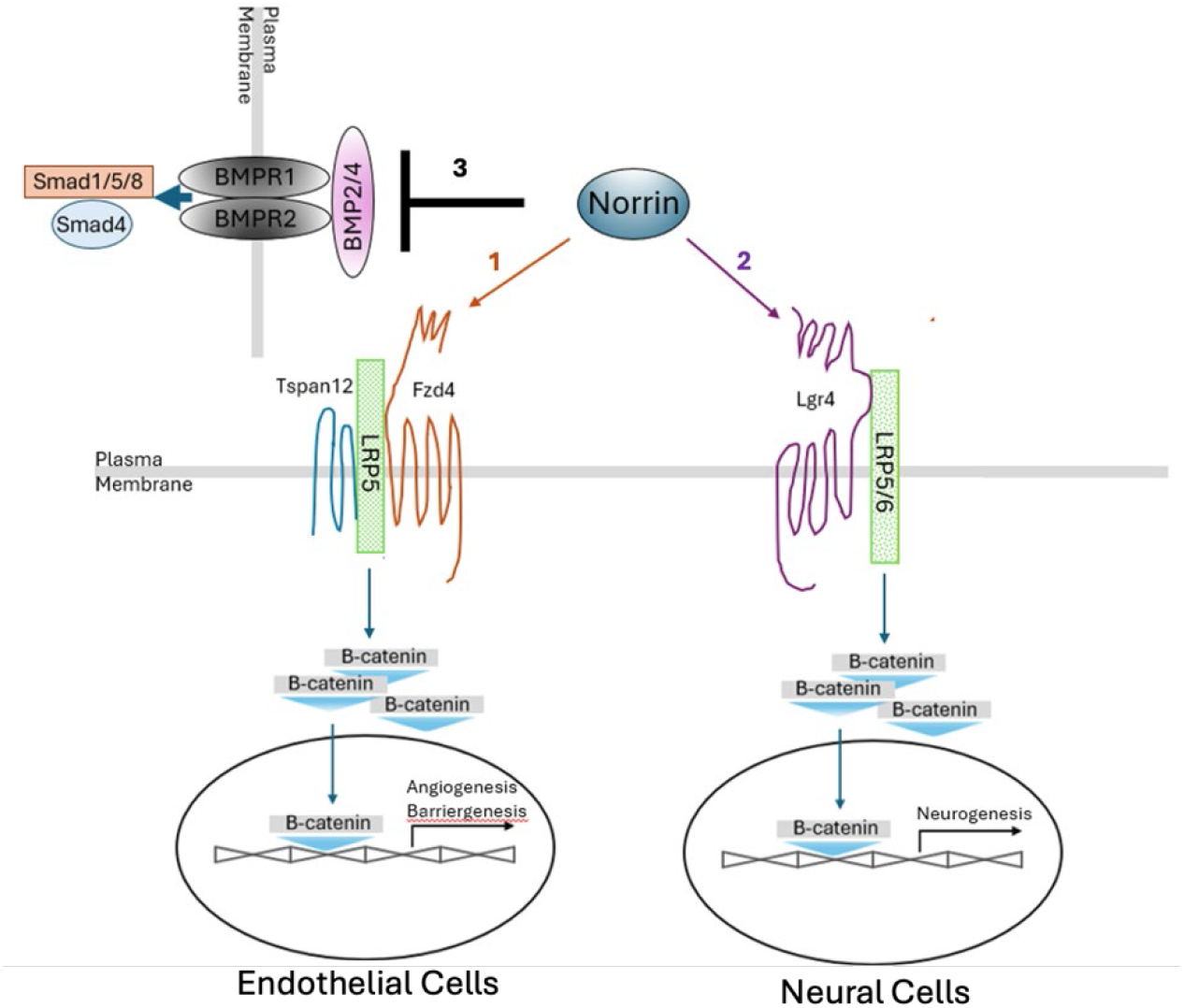
Diagram showing the multifunctionality of norrin. Norrin activates the β-Catenin pathway through both the FZD4 receptor in endothelial cells (1) and the LGR4 receptor in neural cells (2). In addition, norrin inhibits TGF-beta and BMP-2/4 signaling (3).

## Materials and Methods

### Receptor Binding ELISA

An in vitro ELISA was created to assess Noregen (& norrin) potency by its ability to bind to the Frizzled-4 (FZD4) receptor. Briefly, a rhFrizzled-4/FC Chimera (R&D Systems, Minneapolis MN; 5847-FZ-050) was coated onto the surface of a 96-well plate overnight. The next day, the plate was washed with buffered surfactant (R&D Systems # WA126). The surfaces were then blocked casein in PBS and a serial dilution of the test Noregen (0.8-500 ng/mL) was made in the blocking agent. Each of the 7-9 concentrations of Noregen was added to duplicate or triplicate wells of the plate and allowed to incubate for 2 hours. After washing away unbound Noregen, a biotinylated norrin antibody (R&D Systems; BAF3014) was added to the wells and allowed to incubate for 2 hours at room temperature. Following incubation with the antibody, the wells were washed and incubated with Streptavidin-HRP (R&D Systems; DY998), followed by more washing. TMB substrate (R&D Systems, DY999) was added to the wells, the plate stored in the dark and color development was stopped by the addition of 2N Sulfuric Acid after 20 minutes. The absorbance at 450 nm was read on a Bio-Tek Epoch 2 Plate Reader, with data analyzed by four-parameter logistic (4-PL) curve fitting to determine the half maximal effective concentration (EC_50_). Statistical significance between Noregen and norrin groups was determined by two-way ANOVA followed by Šídák’s multiple comparison test.

### TCF/LEF and Smad reporter assays

β-Catenin Reporter assays were conducted by transfecting HEK-293 cells with a TCF/LEF Reporter plasmid (Qiagen Cignal Reporter TCF/LEF, CCS-018L). The Reporter assay is a mixture of inducible TCF/LEF-responsive construct expressing firefly luciferase and a construct constitutively expressing renilla luciferase.

HEK-293 cells (ATCC, Manassas, VA) were cultured using Advanced MEM media (Gibco, Cat # 12492013) containing 5% fetal bovine serum (FBS), 1X Glutamax and Pen Strep. HEK-293 cells were seeded into 24-well plates, grown until 70-90% confluency and then transfected with the Reporter and receptor plasmids using a Lipofectamine 3000 transfection kit Kit (Thermo Fisher Scientific, Waltham, MA). For Noregen dose response assays, 50 ng of pCMV6-XL6-FZD4, 100 ng of pCMV6-XL6-LRP5 & 100 ng of TCF/LEF Reporter plasmids were transfected per well. For evaluation of LGR4, 50 ng of pCMV6-XL4-LGR4 or pCMV6-XL6-EMPTY were also transfected. The receptor and empty vector plasmids were obtained from Origene (Rockville, MD). OptiMEM media that contained no serum or antibiotics was used to prepare transfections. After 5 hours, the media was replaced with Advanced MEM media with 5% serum and antibiotics, and the plate was returned to the incubator.

At 24-hours post transfection, media in wells was replaced with media containing Noregen and incubated for an additional 24 hours. For Noregen dose response assays, eleven dose levels were evaluated using triplicate wells per dose level. For LGR4 assays, three Noregen dose levels (5, 25 & 100 ng/mL) using triplicate wells per dose level were used.

After 24 hours, the HEK-293 cells were washed with PBS, lysed and assayed for firefly luciferase and renilla luciferase activity using the Dual-Luciferase® Reporter Assay System (Promega, E1910) and a 20/20 Luminometer with dual injection (Promega, Madison, WI). Firefly luciferase readings were normalized to the renilla luciferase readings to correct for transfection efficiency and generate relative luciferase units (RLU). Replicate counts ratios (firefly/renilla luciferase) were averaged and four-parameter logistic (4-PL) curve fitting was used to determine the half maximal effective concentration (EC_50_). Statistical significance between each treatment level and the control group was determined by one-way ANOVA followed by Dunnett’s multiple comparison test for the dose response analysis. Statistical significance between LGR4 and control group was determined by two-way ANOVA followed by Šídák’s multiple comparison test.

SMAD reporter assays were conducted by transfecting HEK-293 cells using a Cignal SMAD Reporter kit (Qiagen, CCS-017L). The kit contains a mixture of TGFß-1-responsive firefly luciferase reporter & CMV-responsive renilla luciferase reporter for normalization. Transfection procedures were the same as those used for the β-Catenin Reporter assays except for the use of the SMAD Reporter instead of the TCF/LEF Reporter. Five hours after transfections, the media was replaced with media containing 20 ng/mL of TGF-β1 (R&D Systems, Minneapolis, MN) alone or in combination with Noregen (5, 50 or 500 ng/mL) and cultured for an additional 24 hours prior to cell lysate collection and luciferase readings as described previously. Triplicate counts ratios (RLUs) per group were averaged and results were reported relative to the untreated group. Statistical significance between each treatment level and the control group was determined by one-way ANOVA followed by Dunnett’s multiple comparison test.

### Quantitative PCR analysis of Human Microvascular Retinal Endothelial Cell (HRMEC) Gene Expression

Primary HMRECs were obtained from Cell Systems (Kirkland, WA). They were seeded into six well plates that had been pre-coated with Attachment Factor (Cell Systems, Kirkland, WA) and grown to confluence using fully supplemented EndoGRO-MV media (Millipore, Burlington, MA), a low serum media that does not contain VEGF. Supplements included EndoGRO-LS Supplement (0.2%), *rh*EGF (5ng/mL), L-glutamine (10mM), heparin sulfate (0.75 U/mL), ascorbic acid, (50 µg/mL), FBS (5%) and hydrocortisone hemisuccinate (1 µg/mL). The cells were weaned to media containing no hydrocortisone hemisuccinate prior to stimulation with Noregen or *rh*norrin (R&D Systems, Minneapolis MN;3014-NR). Concentrations ranged from 2-2000 ng/mL, and, after a 24-hour duration, the cells were trypsinized and collected for RNA isolation.

Total RNA was isolated using the Monarch Total RNA Miniprep kit (NEB, Ipswich, MA; T2010) with the optional On-column DNase I treatment to remove residual DNA, according to the kit instructions. First-strand cDNA was synthesized by reverse transcribing 1 µg of total RNA using the LunaScript RT Super Mix Kit (NEB, Ipswich, MA; E3010L). The reaction conditions were according to the manufacturer’s instructions: 25ºC for 2 min, 55ºC for 10 min, and 95ºC for 1 min. qPCR was performed using a duplex reaction format with FAM-labeled probe/primer pairs for the gene of interest and VIC-labeled probe/primer-limited pairs for TBP (Tata-Binding Protein) as the normalizer gene. For real-time PCR reactions, sample first-strand cDNA was diluted 5-fold with deionized water and 2 µL added to 18 µL of master mix for 20 µL PCR reactions. Triplicate reactions were run for each sample using the Luna Universal Probe qPCR 2x Master Mix with Rox reference dye (NEB, Ipswich, MA; M3004). Reactions were performed on an AriaMx Real-time PCR System (Agilent, Santa Clara, CA) and the comparative method used to determine the relative gene expression. Each replicate reaction was internally normalized relative to endogenous TBP gene expression. The specific assay probe sets used for gene expression analysis are listed in Table 1. Statistical significance between treatment and control groups was determined using a one-way ANOVA test followed by Dunnett’s multiple comparison test.

**Table 1.** Taqman gene expression probes for HRMEC gene expression.

| Gene | Probe Set # | Spans Exons |
| --- | --- | --- |
| *PLVAP* | Hs00229941_m1, FAM | 3-4 |
| *AXIN2* | Hs00610344_m1, FAM | 4-5 |
| *TBP* | Hs00427620_m1, VIC-PL | 3-4 |

### ECIS analysis of HRMEC proliferation and barrier resistance

Human Retinal Microvascular Endothelial Cells (HRMEC) were sourced from a 26-year-old human male and used up to the 8th passage (Cell Systems, Kirkland, WA). The cells were cultured using EGM Endothelial Cell Growth Medium Bullet Kit (Lonza, Basel, Switzerland) containing 0.4% Bovine Brain Extract, 10 ng/ml EGF, 0.1% Ascorbic Acid, GA-1000, FBS and 100nM Hydrocortisone (HC). The level of FBS was 5% during cell attachment and reduced to 1% during experiments. Prior to cell culture, 96-well ECIS plates (96W20idf PET, Applied BioPhysics, Troy, NY) were treated with 10 mM cysteine, for 15-minute treatment, the solution was aspirated and two washes with distilled water were completed. Wells were then treated with cell attachment factor solution for 15 minutes (Cell Systems, Kirkland, WA) to prepare for cell seeding. For cell proliferation experiments, HRMECs were seeded into the prepared 96-well ECIS plates at a density of three thousand cells per well using 0.2 mL media per well. After allowing the plates to sit at room temperature for 20 minutes, the plates were placed in the incubator (37 °C, 5% CO_2_) for attachment. After 6 hours, the seeding media containing 5% FBS was aspirated and replaced with 0.2 mL of media containing 1 % FBS with Noregen or norrin at 20 ng/ml or 500 ng/ml. The plates were then inserted into the plate holder of the Z-Theta model ECIS system (Applied Biosystems, Troy, NY) and data collection started. The time course measurements were set to measure capacitance and resistance at multiple frequencies every 1800 seconds (30 minutes). Data collection lasted for 45 hours, which was 51 hours post-seed. The capacitance at 32,000 Hertz was used to assess cell proliferation and the capacitance of 8 wells per group were averaged. Statistical significance between the groups was determined using a two-way ANOVA test followed by Tukey’s multiple comparison test.

For experiments measuring the barrier recovery, 96-well ECIS plates were prepared for cell seeding in the similar manner. A higher number of HRMECs per well (20,000) were seeded using EGM Endothelial Cell Growth Medium Bullet Kit containing 5% FBS and 1 µM HC and the cells were placed in the incubator overnight. The next day the media was replaced with media containing 1% FBS and 1 µM HC, the plate was placed in the ECIS system and resistance measurements were collected as previously stated. After 16 hours, the media in the wells was replaced with 1. Media alone, 2. Media containing 50 ng/ml Vascular Endothelial Growth Factor (VEGF165, R&D Systems, Minneapolis,MN) or 3. Media containing VEGF165 in combination with 40 ng/ml of Noregen. Six replicate wells were treated per group. The resistance measurements at 4000 HZ were normalized to initial values and replicates were then averaged at each time point over 80 hours. Statistical significance between the VEGF alone group and the VEGF & Noregen co-treatment group was determined using a two-way ANOVA test followed by Tukey’s multiple comparison test.

### Oxygen Induced Retinopathy Model

An Oxygen Induced Retinopathy Model (OIR) first described by Smith et al was employed.^8,9^ C57BL/6J pregnant mice were purchased from Charles River. Upon giving birth, the dams and pups were put into a 75% oxygen chamber on postnatal day (P) 7 for 5 consecutive days. The dams were rested by taking them out of the high oxygen chamber for 1 hour each day. On P12 the pups and dams were returned to room air. After removal from the high oxygen chamber the pups received intravitreal injections (1 µL) of either Noregen, norrin or vehicle on P13 (or P14 for Noregen, 20 ng cohort) in the right eyes (OD) only using a 34-gauge beveled needle (WPI, Sarasota, FL). Four sets of injections were made in the pups after removal from oxygen. Cohort A; Noregen-2 ng (n=8) & Vehicle (n=8) Cohort B; Noregen-20 ng (n= 6) & vehicle (n=7), Cohort C; Noregen-200 ng (n=9) & vehicle (n=9) and Cohort D; Norrin-200 ng (n=11) & vehicle (n=10). The vehicle was PBS in all cohorts except for Cohort B where it consisted of Noregen formulation buffer [20mM Histidine, 250mM Sucrose, 0.01% PS-20, pH 5.5]. The left eyes (OS) served as internal controls since mouse weight has been shown to affect the timing and severity of the retinopathy.^9^ On P17 mice were sacrificed, eyes were fixed in 4% PFA for 1 hour, washed with PBS and the retinas were isolated. The retinal vessels were stained with Isolectin B4 Alexa 495 (ThermoFisher Waltham, MA) at room temperature overnight. The stained retinas were then washed 3-4 times for 15 minutes with PBS and mounted onto slides. Images of the whole mounted retinas and 3D image stacks were obtained on a Ziess LSM900 AIRYSCAN confocal microscope. To calculate the Avascular Area (AVA), we used GIMP (GNU Image Manipulation Program) software to measure the pixels in the AVA and divide by the pixels in the total retina. To determine the normalized AVA, the percentage of AVA from OD eyes was divided by that of fellow un-injected OS eyes. The percent reduction in AVA was then calculated by subtracting the normalized AVA from one. Significance after normalization was determined using a one-tailed Student’s t-test to compare groups treated with Noregen or norrin to vehicle groups in each cohort.

### Immunofluorescence

Primary HMRECs were seeded and grown to confluence in 24-well chamber slides that had been pre-coated with Attachment factor. Fully supplemented EndoGRO-MV media containing 5% FBS and 2 µM hydrocortisone hemisuccinate (HC) culture media was used to establish tight junctions and then the serum and HC were reduced to 1% and 0.1 µM, respectively, one day prior to treatments. Cells were treated with human Vascular Endothelial Growth Factor (VEGF165) (50 ng/mL), bone morphogenetic protein 2 (BMP-2) (50 ng/mL) or Transforming Growth Factor beta 1 (TGF-ß1) (20 ng/mL) alone or in combination with Noregen (100 ng/mL for VEGF_165_ and 20 ng/mL for BMP-2 & TGF-ß1). All additives were purchased from R&D Systems (Minneapolis, MN). After 24 hours, the cells were fixed for 10 minutes with 4% paraformaldehyde. The cells were washed with PBS & permeabilized with 0.1% Triton-X100 for 15 minutes followed by blocking with 3% BSA for 45 minutes. For detection, cells were incubated overnight at 4°C with rabbit polyclonal primary antibody recognizing the tight junction protein ZO-1 (ThermoFisher, 40-2200), followed by washing and incubation for 1 hour with Goat Anti-Rabbit IgG (H+L) Alexa Fluor® 488 secondary antibody (ThermoFisher, A-32731). After washing with PBS, the coverslips were mounted using Mounting media containing DAPI. Images were taken using a Cytation 5 Cell Imaging Multi-Mode Reader (Agilent, Winooski, VT).

### LGR4 immunostain of retinal cross-sections

The eyes from adult C57BL/6J mice were fixed in Davison’s fixative and then embedded in paraffin. 5 µm sections were cut and mounted onto slides for immunostaining. Slides were dewaxed and rehydrated by immersion in xylene followed by decreasing concentrations of alcohol solution and lastly water. Citrate buffer, pH 6.0 was used as an antigen retrieval buffer. Endogenous peroxidase was quenched with 3% peroxide for 30 minutes and after rinsing with PBS the slides were blocked with 2% goat serum containing 0.2% TritonX-100 in PBS for 30 minutes. Antibody specific for leucine-rich repeat GPR 4 (LGR4) (Novus, Centennial, CO) was diluted 1:500 with blocking solution, pipetted onto slides and incubated overnight at room temperature. The following day, an immPACT SG Substrate Kit (Vector Labs, Newark, CA) was used to enzymatically stain LGR4 and nuclear Fast Red was used as a counter stain. The slides were digitized using a 20X objective lens and SL120 Virtual Microscopy slide scanner (Olympus, Center Valley, PA).

### Statistics

Graphs represent mean+/-SD except where indicated. Values are means +/-SD (or SEM); *P<0.05, **P<0.01, ***P<0.001 & ****P<0.0001. ANOVA and Student’s t-test were performed using GraphPad Prism version 10.4.2 (GraphPad Software, Boston, MA).

## Results

### Confirmation of GMP scalable manufacturing methods

Following creation of the engineered recombinant norrin-derived growth factor Noregen, peptide mapping using trypsin digest confirmed that its primary sequence is correct. Predictive conformational protein tertiary structure modeling confirmed that Noregen and norrin share the same three-dimensional structure. Intact LC/ES-MS revealed a dimer with the predicted molecular weight and correct number of disulfide bridges was present at 98%. Sedimentation velocity analytical ultracentrifugation (SV-AUC) was used to determine the main species of the Noregen drug substance in its native state. Lower and higher [order] molecular species represented a very small proportion with the main dimer peak being 99.9% of total content.

To confirm Noregen’s potency, multiple batches were tested in a previously developed and qualified receptor binding ELISA. An average EC_50_ of 22 ng/mL (+/-4 ng/mL) was determined upon testing three of the most recent Noregen batches. An example 4-PL curve fit (Fig. 2a) comparing Noregen to human recombinant norrin (R&D Systems, Minneapolis, MN) demonstrates that Noregen has an equal, if not superior, FZD4 receptor binding efficacy compared to that of norrin. The determined EC_50_s were 19 ng/mL and 34 ng/mL for Noregen and norrin respectively.

**Figure 2.**
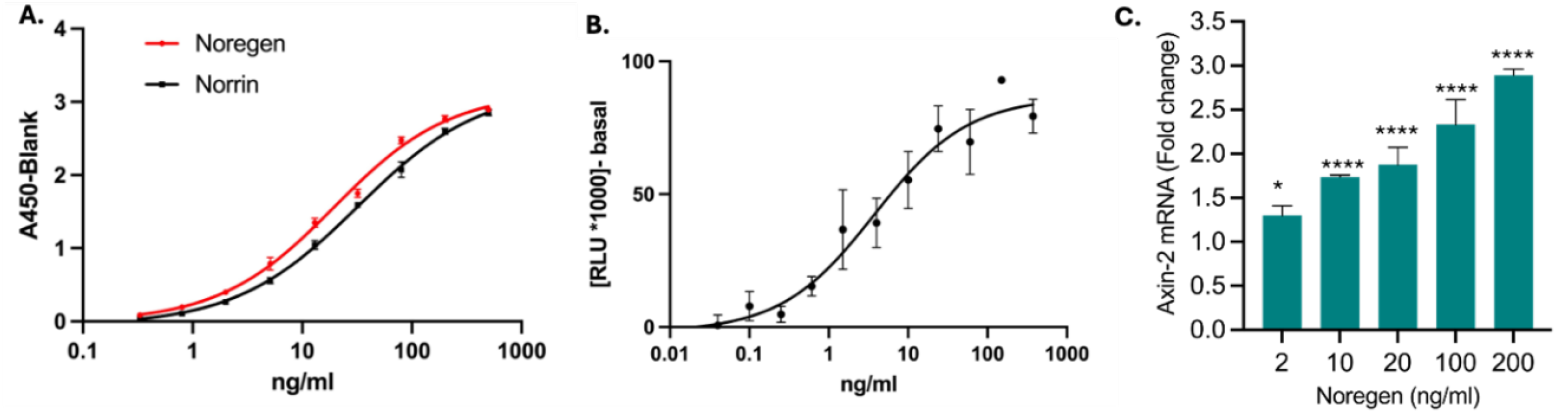
Confirmation of GMP scalable manufacturing. (A) Noregen FZD4 receptor binding activity compared to Norrin, EC_50_ 19 ng/mL & 34 ng/mL respectively. (B) Noregen dose response curve for TCF/LEF reporter activation in HEK-293 cells (EC_50_ 11 ng/mL). (C) Noregen induced Axin-2 gene expression in HRMECs. Values are means +/-SD; *P<0.05, **P<0.01, ***P<0.001 & ****P<0.0001.

The best characterized norrin-Wnt/β-Catenin signaling pathway functions via complexing of the Frizzled-4 receptor (FZD4) with a co-receptor, low-density lipoprotein receptor-related protein 5 (LRP5). Upon binding to these cell surface receptors, intracellular β-catenin is freed and translocated to the nucleus where it binds to TCF/LEF transcription factors and activates target gene expression. We evaluated Noregen activation of β-Catenin signaling in HEK-293 cells that were transfected with vectors expressing FZD4, LRP5 and firefly luciferase under the control of TCF/LEF transcription binding elements. Six assays were performed, each with 3 replicates per dose level, to evaluate 3 lots of Noregen (Fig. 2b) where the average EC_50_ was 11 ng/mL (+/-7 ng/mL). This compares favorably to 3 assays done with human norrin with an average EC_50_ of 36 ng/mL (+/-17 ng/mL).

Another assay suitable for evaluation of norrin-Wnt/β-Catenin signaling activation is Axin-2 mRNA gene expression. In HRMECs, Noregen increases AXIN2 expression in a dose dependent fashion (Fig. 2c), and this response is similar to human norrin treatment (data not shown).

### Retinal endothelial cell tight junctions and anti-permeability

Plasmalemma vesicle-associated protein (PLVAP) plays a major role in cellular transcytosis and is a marker of fenestrated endothelium.^10,11^ High levels are associated with vascular endothelial cell growth factor (VEGF)-induced ischemic retinopathies, such as diabetic retinopathy, retinal vein occlusion, familial exudative vitreoretinopathy (FEVR) and pathologic angiogenesis as seen in age-related macular degeneration and neovascular proliferation.^10,12^ For this reason we looked at the gene expression of PLVAP in HRMECs treated with Noregen or norrin (Fig. 3a). Incubation with Noregen for 24 hours resulted in a dose dependent decrease in PLVAP expression. A Noregen dose of 20 ng/mL resulted in a 2-fold decrease in PLVAP expression, which is comparable to a 2-fold reduction seen with norrin treatment at ten times the dose (200 ng/mL).

**Figure 3.**
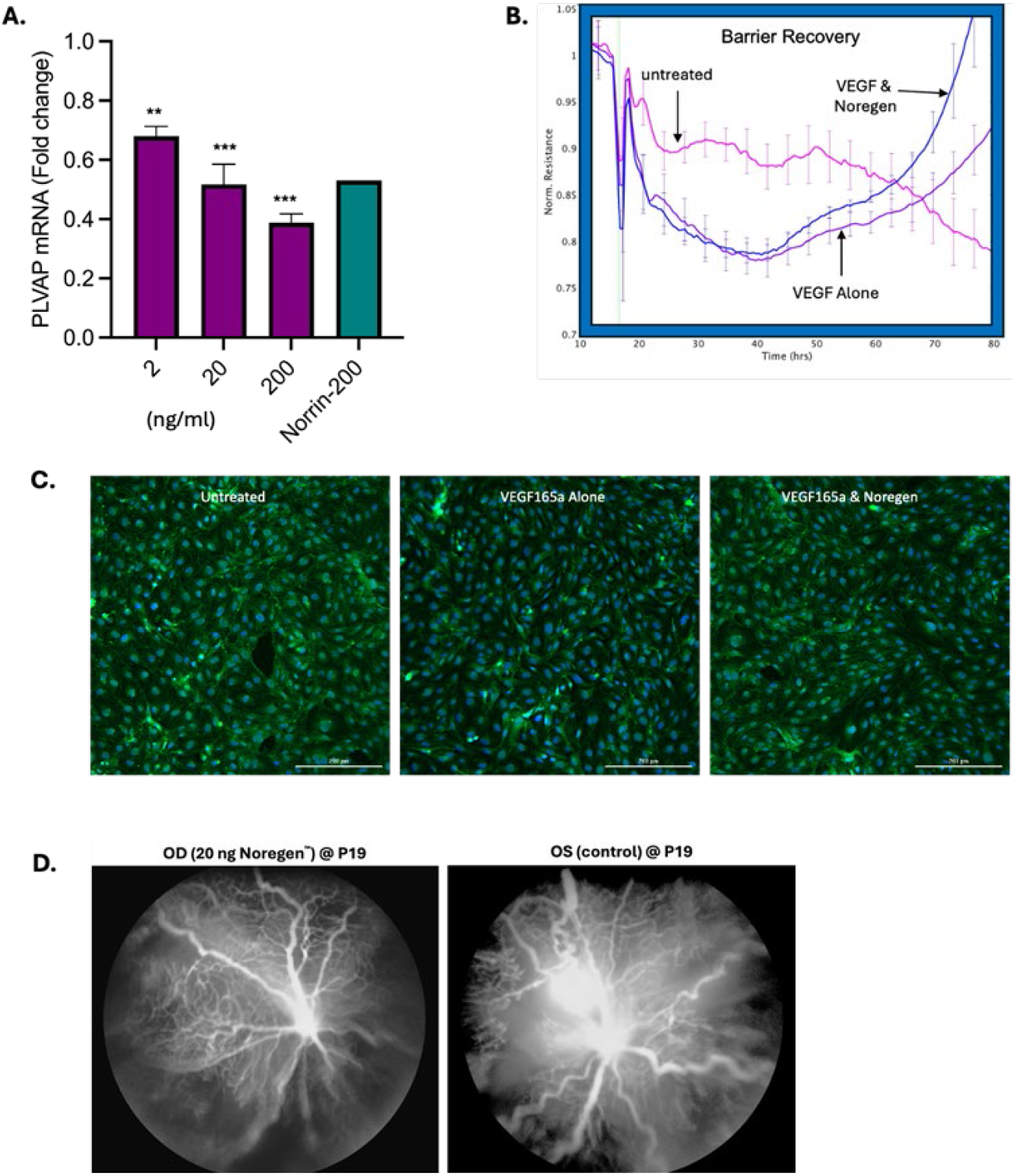
Noregen induced Barrier. (A) Noregen decreases PLVAP expression, a marker of fenestrated endothelium, in HRMECs. Values are means +/-SD; *P<0.05, **P<0.01, ***P<0.001. (B) Noregen induced resistance recovery in HRMEC monolayer depleted by VEGF165. Significantly higher normalized resistance in the Noregen co-treated group compared to group treated with VEGF165 alone (p<0.05 from 52 hours to end). (C) Noregen restoration of tight junction, ZO-1, at HRMEC membrane after depletion by VEGF165a. (D) Comparison FA’s in OIR eyes injected with Noregen in right eye on P14 and fellow eye that was un-injected. Images were acquired on P19.

One of the pathological effects of elevated VEGF levels is the loss of functional blood-brain(retina)-barrier. To assess HRMEC barrier recovery after VEGF injury an Electrical Cell-substrate Impedance Sensing (ECIS) Z-Theta System was employed. For this assay, HRMECs were grown to confluence in 96-well ECIS plates and the resistance (R) across the monolayer was monitored over several days. At frequencies lower than 4000 HZ, the impedance is primarily due to resistance of current flowing in the paracellular space between cells. Three test groups were evaluated with 6 replicate wells per group: VEGF165 (50 ng/mL) alone, VEGF165 (50 ng/mL) together with Noregen (40 ng/mL), or media alone (untreated). VEGF165 induced a significant (24%) drop in resistance 24 hours after addition to the culture media and the resistance recovered to untreated levels by 48 hours. Noregen accelerated barrier recovery as can be seen by the significantly higher normalized R values after the initial depletion by VEGF165 (Fig. 3b). A significant difference (<0.05) was reached 52 hours after the start of data collection (36 hours after treatment start) and maintained through the end of the experiment.

VEGF depletes tight junction proteins such as Claudin-5 and ZO-1 located at the boundaries of HRMECs, whereas norrin restores barrier function by translocating these proteins to the cell membranes.^4,13^ We found that co-incubation with Noregen (100 ng/mL) restores ZO-1 in VEGF treated (50 ng/mL) HRMECs (Fig. 3c). In addition, cells treated with VEGF165-a alone were elongated and more disorganized than those treated with a combination of VEGF165-a and Noregen.

The OIR mouse model with *in vivo* imaging highlights the permeability effects of increased VEGF in the setting of ischemic retinopathy. Notably, vessel dilation and tortuosity, vessel permeability and capillary drop-out can be appreciated in this model. *In vivo* live imaging using fluorescein angiography shows that, when compared to fellow eyes, Noregen treated eyes had normalized retinal vascular appearance with straightening of the vessels, decreased vessel permeability, and increased capillary networks (Fig.3d).

### Retinal vascular regeneration

We and others have shown that norrin promotes the revascularization of the retina after vaso-obliteration in the mouse OIR model.^14,15^ In this study, we looked at Noregen induced proliferation of HRMECs by measuring capacitance with an ECIS Z-Theta system. Specifically, a low number of cells per well were seeded into a 96-well ECIS plate that contains gold electrodes at the bottom of each well. As the cells grow, the capacitance (C), which is indirectly proportional to the number of cells on the electrode, is measured. A lower C value indicates cell proliferation. We compared the C in cells treated with either Noregen or norrin at two levels, 20 ng/mL and 500 ng/mL (Fig. 4a-c). Both Noregen and norrin significantly decreased capacitance compared to the untreated cells, indicating an increase in proliferation at the high dose level, 500 ng/mL (p<0.01 from 5 hours to 28 hours). Noregen, however, also significantly decreased the capacitance at the 20 ng/mL dose (p< 0.01at every time point until end). This confirms that Noregen promotes endothelial cell proliferation equal or superior to norrin.

**Figure 4.**
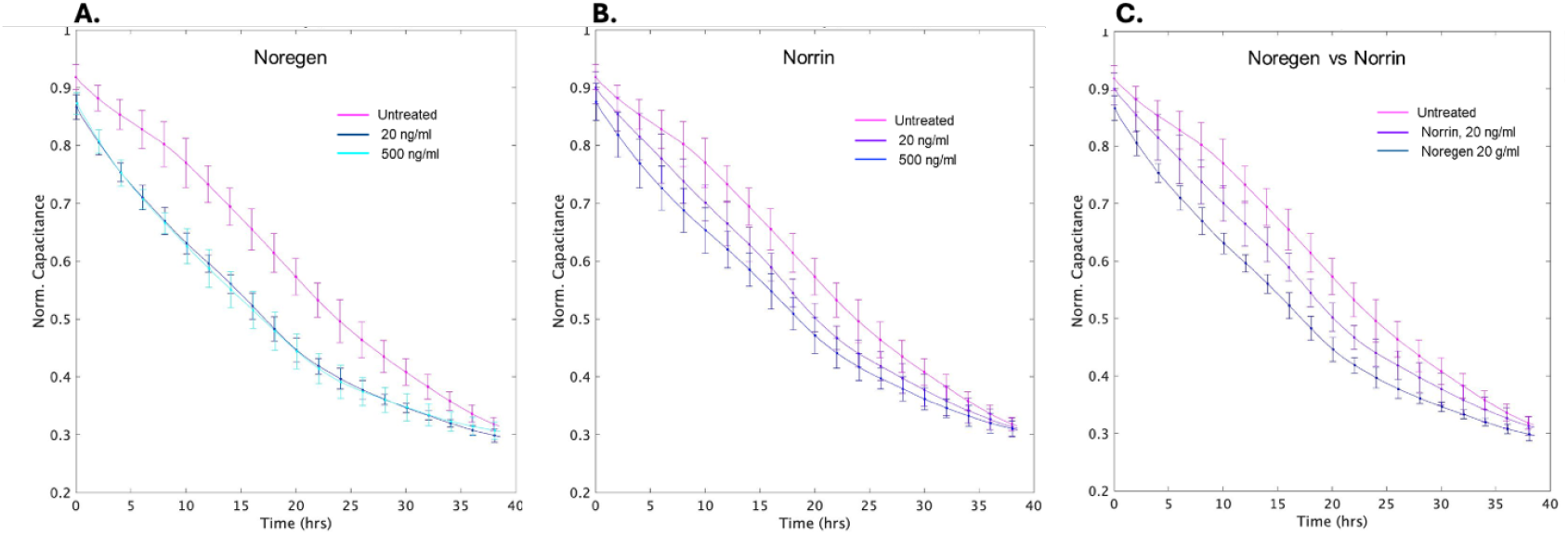
Noregen induced proliferation. (A) Noregen significantly increases HRMEC proliferation as seen by a drop in capacitance when treated at 20 and 500 ng/mL. (B) Norrin increases proliferation to a similar extent as Noregen at the higher 500 ng/mL treatment level, but to a lesser extent at 20 ng/mL. (C) Comparison of Noregen and norrin treatments at 20 ng/mL. An earlier drop in capacitance indicates faster growth in Noregen treated cells.

We have previously shown that an intravitreal injection of recombinant norrin into mouse OIR eyes on P14 (after ischemic pathology is induced) and evaluated at P17 (height of pathologic changes), significantly rescues the vaso-obliterated retina, promotes the growth of non-fenestrated retinal vessels and reduces neovascular tufts (ischemic retinopathy).^14^ Furthermore, the results demonstrated the effect that mouse pup weight has upon avascular and neovascular areas, as had been previously illustrated by Smith and colleagues.^9^ To overcome this variability due to post-natal weight here, we made injections in only one eye (OD) and normalized results to the fellow un-injected eyes (OS). The original study was repeated using logarithmic dose escalation for Noregen (2 ng, 20 ng, 200 ng) and repeating norrin (200 ng) compared to vehicle injections. The 200 ng doses of norrin and Noregen were comparable to each other and confirmed the results from the original study with intravitreal norrin treatment in the OIR model (Fig. 5a). The 200 ng dose of Noregen replicated the norrin results with a 40% reduction in normalized retinal avascular area and reached statistical significance compared to vehicle treated eyes (p=0.006). The 20ng dose also showed a reduction in avascular retinal area with a trend towards significance (p=0.08) compared to vehicle treated eyes and was significant compared to un-injected internal control eyes (p=0.05). The 2ng Noregen dose was equivalent to vehicle treated eyes. Figure 5b contains a compilation of the results from all cohorts with averaged vehicle results. Fig. 5c presents example images of whole mounted retinas with highlighted avascular areas.

**Figure 5.**
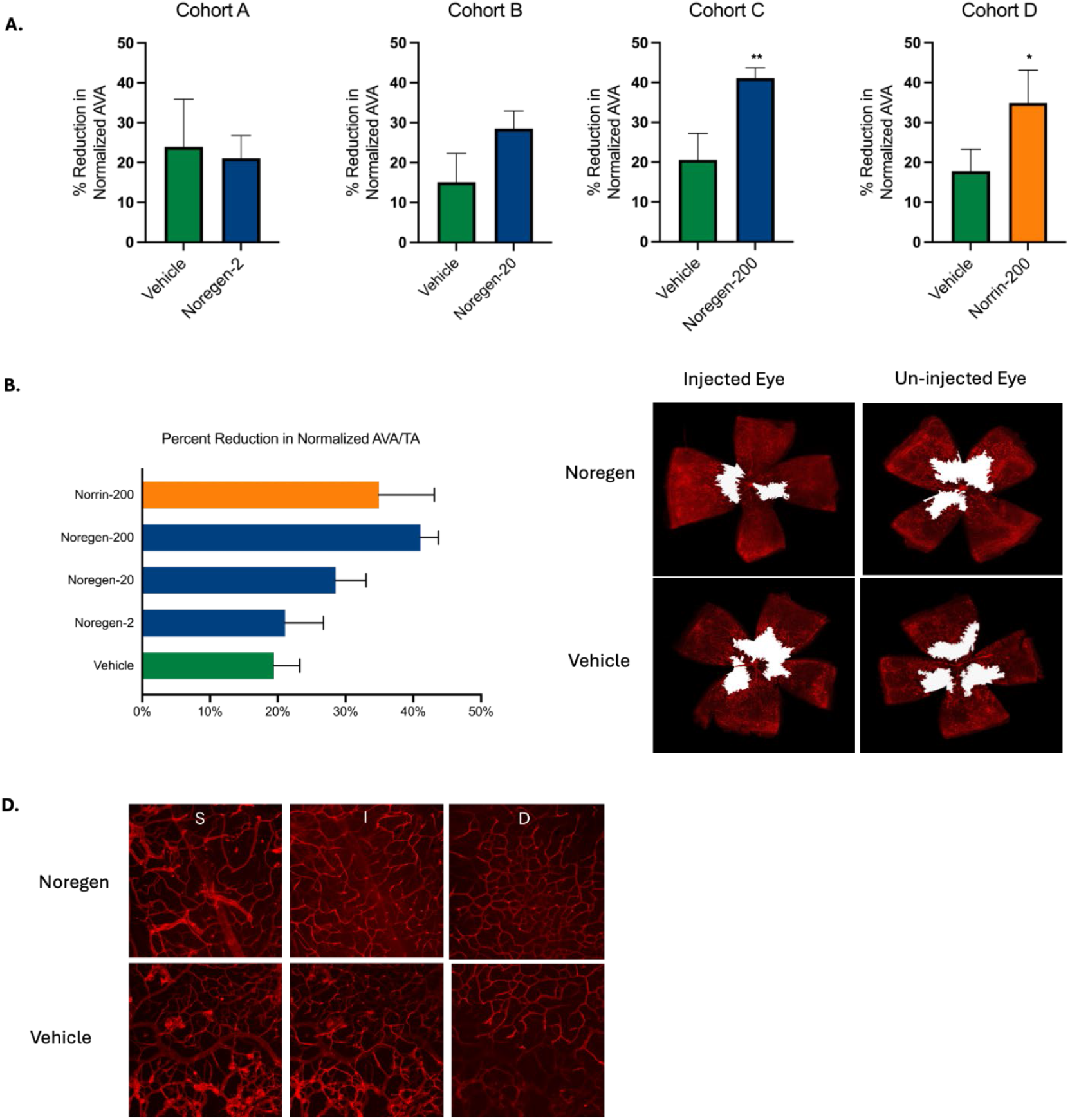
Noregen retinal vascular recovery in OIR model. (A) Reduction in normalized avascular area [AVA to Total Area ratio] in Noregen & norrin injected eyes. The [AVA to Total Area ratio] in injected eyes was normalized to fellow un-injected eyes. Noregen levels were 2, 20 or 200 ng. Values are means +/-SEM; *P<0.05, **P<0.01. (B) Dose dependent reduction in normalized avascular area seen when graphing in comparison to averaged Vehicle from combined cohorts. (C) Example retinal flatmount images with AVA highlighted in white. (D) Confocal images of superficial (S), intermediate (I) & deep (D) vascular beds in Noregen(200 ng/eye) and vehicle injected OIR eyes.

It is well documented that the OIR mouse retina will spontaneously recover the superficial capillary plexus by P29; however, the intermediate and deep plexi do not recover.^9^ Therefore, we used confocal microscopy to image the superficial, intermediate and deep capillary plexi. The whole mount retinas showed improved superficial capillary regeneration in treated eyes compared to untreated eyes, and confocal imaging showed that Noregen-treated eyes additionally improved intermediate and deep capillary growth (Fig. 5c and d).

### Activation of neurogenic Wnt signaling

Multiple *in vivo* animal studies have shown that the introduction of norrin in the eye, either via intravitreal injection or expression from transfected host cells, is neuroprotective and neurogenic. ^16–19^ Norrin has been shown to be a key regulator of neuroprotection and neurogenesis in Wnt-driven and Wnt-independent pathways. In addition to Wnt signaling via the FZD4 receptor, norrin activates another β-Catenin signaling pathway via the LGR4 receptor, which is associated with neurogenesis.^3,20^ FZD4 and LGR4 cannot simultaneously bind norrin due to steric interference, supporting that norrin-LGR4 binding is another Wnt signaling pathway which is independent from FZD4 binding.^20^ LGR4, 5 and 6 were first identified as key receptors in the activation of progenitor stem cells in the crypt cells of the intestine. LGR4 is expressed in the inner plexiform layer and retinal ganglion cells of the retina (Fig. 6a), activating neurogenesis within the retina. In this study, LGR4 enhancement of Noregen-induced β-Catenin activity was evaluated with a TCF/LEF dual luciferase assay. HEK293 cells were transfected with a TCF/LEF Reporter and either LGR4 (50 ng) or empty vector. Transfected cells were treated for 24 hours with Noregen (0, 5, 25 or 100 ng/mL). The Relative Luciferase Units (RLU) were higher in the LGR4 transfected cells at every concentration of Noregen with a maximum response seen with 25 ng/mL Noregen (Fig. 6b). LGR4 enhanced the signaling by a factor of 1.9 at the 5 ng/mL (p=0.02) and a factor 1.5 at the 25 ng/mL (p=0.02) Noregen dose levels. This data confirms that Noregen activates LGR4-mediated Wnt signaling similar to human norrin.

**Figure 6.**
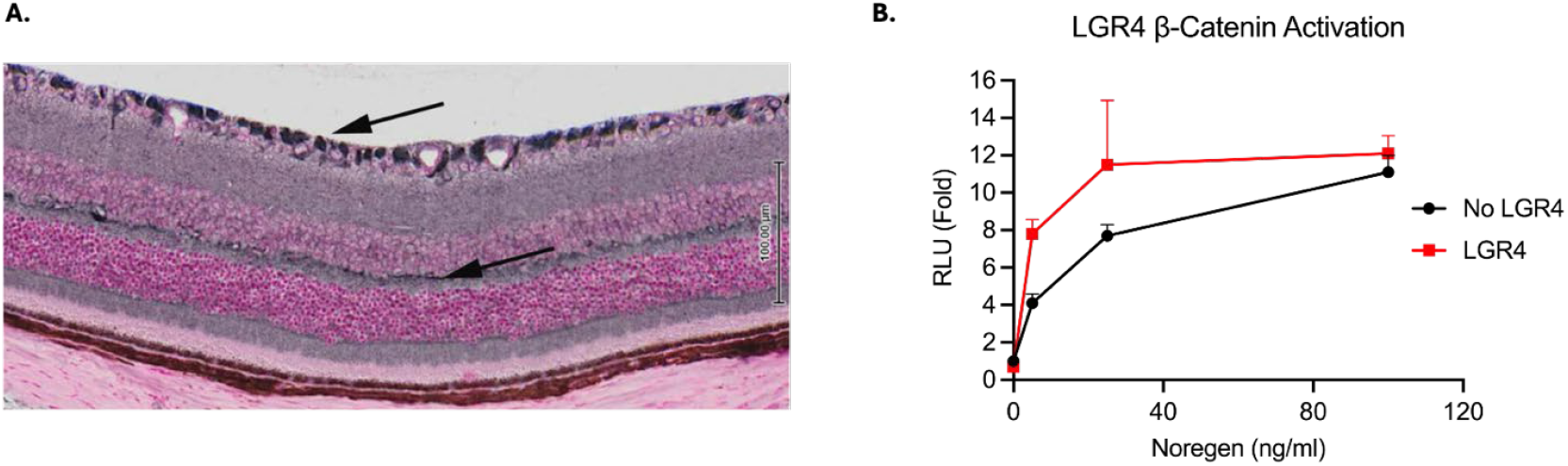
Noregen ß-Catenin Activation via LGR4 receptor. (A) Localization of LGR4 in RGC and IPL layers of adult C57BL-J mouse retinas demonstrated using enzymatic immunostaining. (B) TCF/LEF Luciferase activation after stimulation by Noregen in HEK293 cells that had been transfected with LGR4 or empty vector. A significant enhancement in signal is seen at 5 and 25 ng/mL of Noregen when LGR4 is present (p= 0.02 at both Noregen levels).

### Anti-fibrosis

Retinal fibrosis is a common complication of ischemic retinopathies, such as diabetic retinopathy and retinal vein occlusion.^21^ Elevated expression of transforming growth factor beta 1 (TGF-β1) has been implicated in retinal fibrosis in proliferative retinopathy^22–24^ via increased expression of extra cellular matrix proteins (ECM), resulting in tractional retinal detachments, a blinding end-stage event in all ischemic retinopathies.^25^ Two other members of the TGF-β superfamily, bone morphogenic protein (BMP)-2 and BMP-4 are also predicted to play a role in diabetic retinopathy (Fig. 7a).^26,27^ Norrin inhibits TGF-β1 signaling by induction of SMAD7 expression.^28^ Norrin has also been found to inhibit BMP-4 and BMP-2 signaling in luciferase reporter assays.^3,6^ In this study, we used a SMAD reporter assay to determine activation of the TGF-β1 pathway in HEK293 cells treated with or without Noregen. We found that Noregen decreases basal levels of SMAD signaling in a dose dependent manner, as well as inhibiting TGF-β1 induced SMAD activity (Fig. 7b). Basal SMAD activity levels decreased by 30%, 60% and 70% when incubated with 5, 50 & 500 ng/mL of Noregen respectively. Incubation with TGF-β1 at 20 ng/mL increased SMAD activity by a factor of 4. Interestingly, co-incubation with 5 ng/mL of Noregen increased activity over that of TGF-β1 treated cells. However, co-treatment with Noregen at the 50 ng/mL and 500 ng/mL levels significantly reduced the TGF-β1 activity by 40% (p<0.01). We also show the depletion of ZO-1 tight junction protein at the HRMEC cell membrane caused by incubation with either TGF-β1 or BMP-2 and its barrier restoration by Noregen (Fig. 7c). This supports the conclusion that Noregen prevents breakdown of the blood-retinal-barrier, which is a precursor to fibrosis, and inhibits SMAD-induced ECM production.

**Figure 7.**
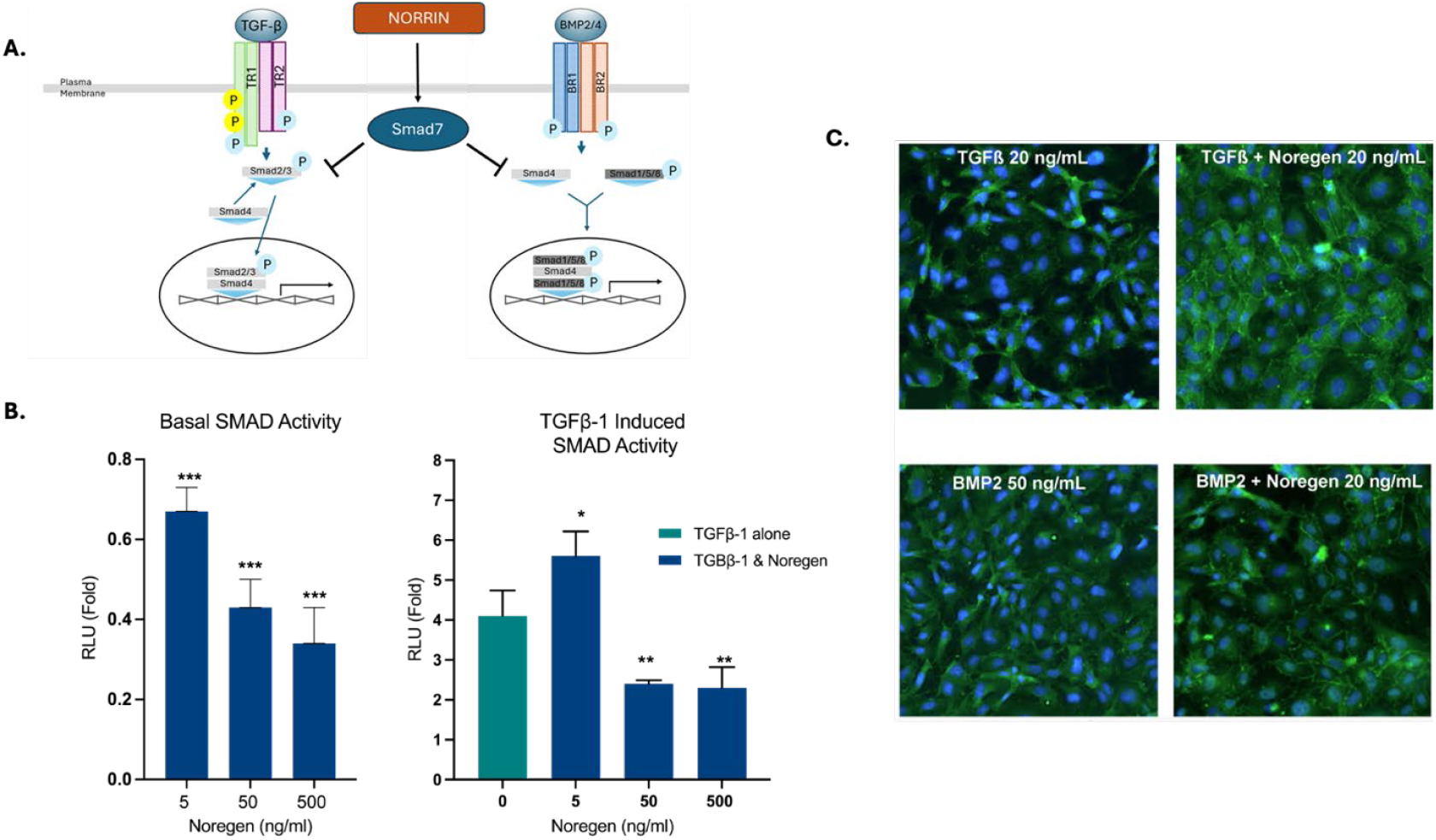
Noregen counters TGF-ß1 and BMP-2 signaling. (A) Diagram illustrating TGF-ß1 and BMP-2 pathways. Norrin inhibits signaling via SMAD7 induction, thereby inhibiting pathway activation. (B) Noregen reduction in basal level and TGF-ß1 induced SMAD reporter activation. Values are means +/-SD; *P<0.05, **P<0.01, ***P<0.001. (C) ZO-1 immunostaining of HRMECs treated with TGF-ß1 or BMP-2 alone (translocation of ZO-1 to cytosol) or in combination with Noregen (restoration of ZO-1 at plasma membrane).

## Discussion

Chronic ischemic retina, both inherited and acquired, results in overproduction of VEGF which causes breakdown of the blood-retina-barrier (macular edema; exudation; hemorrhage), pathologic angiogenesis (neovascularization; proliferative retinopathy), and peri-retinal cellular proliferation and fibrosis (tractional retinal detachment). The current standard of care is blockade of VEGF to repress vascular permeability and neovascularization. However, there is no current treatment to regenerate the retinal vasculature, restore and maintain the blood-retina-barrier, or address the neurosensory loss and progressive fibrosis associated with ischemic retina. The discovery of the novel multi-functional ligand norrin as a key regulator in retinal development, including the growth of non-fenestrated retinal vessels and maintenance of the blood-retina-barrier, led to the preclinical data presented here that demonstrate a recombinant engineered norrin-derived growth factor, Noregen, can be used therapeutically to rescue ischemic retina.

The most well-characterized norrin signaling pathway is FZD4 receptor-mediated canonical Wnt signaling. However, norrin binds FZD4 alone or as the co-receptor complex of FZD4/LRP5/TSPAN12 and activates, through these receptors and complexes, both canonical and non-canonical pathways. Canonical and non-canonical norrin-FZD4 pathways are critical for non-fenestrated vessel growth and the generation and maintenance of the blood-retina-barrier. Canonical signaling is required for tip cell sensitization and non-canonical signaling is required for stalk cell proliferation.^29^ FZD4 canonical and non-canonical signaling contributes to the blood-retina-barrier, and norrin acts as both a canonical and non-canonical ligand in the growth and maintenance of non-fenestrated vessels. Here we present data that confirms that Noregen binds the FZD4 receptor and activates Wnt-β-catenin signaling as well or better than human norrin.

In order to demonstrate that norrin also activates FZD4-mediated canonical and noncanonical signaling, resulting in growth of non-fenestrated retinal vessels, we and others have published on the rescue of OIR in the mouse model^14^ and the prevention of retinopathy.^15^ Additionally, our previous work has shown that treatment with anti-VEGF (aflibercept) worsens retinal avascularity in the same OIR mouse model, resulting in structural and functional deficits.^30^ Importantly, norrin-mediated pathway activation not only promotes vessel revascularization, but also promotes vessel integrity with an appropriate blood-retina-barrier.^4^ Data presented here show that our norrin-derived growth factor also promotes retinal vascular regeneration in the OIR mouse model in a dose-dependent fashion with equal efficacy to human norrin. *In vivo* imaging of OIR mice following Noregen treatment show dramatic decrease in vessel dilation and tortuosity, vessel leakage, and capillary drop-out. Data presented here confirms that Noregen restores barrier function and the blood-retina-barrier, in part by inhibiting transcytosis via PLVAP and translocation of tight junction protein, zonula occludens-1, to the retinal endothelial cell membrane in the same fashion as human norrin.

Another norrin-mediated Wnt pathway receptor is LGR4. LGR4, 5, and 6 are well-characterized receptors of key importance to stem cell activation.^31^ LGR5 mediated Wnt signaling has been shown to be a key receptor in the regeneration of hair cells of the cochlea^32,33^ and norrin binds to LGR5.^3^ Norrin, via LGR4 binding and pathway activation, has been shown to protect damaged retinal neurons and induce retinal progenitor proliferation^20^, and our previous work with norrin demonstrated increased retinal ganglion cell (RGC) number and improved dendrite maturation as well as improved retinal thickness and organization in the OIR mouse model following intravitreal norrin treatment.^34^ Additionally, norrin and LGR6 expression are enriched in astrocyte and glial cells and both modulate neuronal activity, dendritic growth and spine formation.^20^ Wnt signaling via norrin binding LGR4 uses both LRP5 and LRP6 as co-receptors.^3^ Similar to mutually exclusive binding of norrin to FZD4 or LGR4, overlapping binding sites between Tspan12 and LRP6 for norrin binding results in competition between FZD4 and LGR4 pathway activation.^20^ In the retina, LGR4 is expressed in RGCs and the inner plexiform layer. Our recombinant norrin-derived growth factor is able to bind and activate both FZD4- and LGR4-mediated Wnt signaling pathways.

The end-stage of ischemic retinopathies such as diabetic retinopathy and retinal vein occlusion is the production of extracellular matrix (ECM) and fibrosis resulting in tractional retinal detachment, a blinding event. Norrin inhibits extracellular matrix expression via multiple pathways. Transforming growth factors of the beta-class (TGF-ß) stimulate extracellular matrix synthesis and have been implicated in embryogenesis, wound healing and fibroproliferative responses to tissue injury.^24,25^ Norrin, as a cysteine-knot protein, acts as do other proteins in this family via direct interaction with other proteins within the family, but also by altering the expression of extracellular proteins via SMAD signaling. Similar to VEGF and DAN proteins^35^, norrin relies on the appropriate tertiary structure of its dimer formation to directly interact with other proteins.^6^ Norrin is a direct antagonist of TGF-β1and BMP2.^36^ Elevated expression of both TGF-β1 and BMP-2 have been implicated in retinal fibrosis in proliferative retinopathy via increased expression of extra cellular matrix proteins. Incubation of HRMECs with BMP-2 or TGF-ß1 results in increased cell permeability^37^ and a model of corneal fibrosis results in increased extracellular matrix with cell contraction.^38^ Seitz *et al* (IOVS 2018) demonstrated norrin-induced inhibition of the TGF-β1 target gene PAI-1 in HRMECs.^36^ In the presence of high glucose, BMP-2 upregulates BMP receptors, thereby activating the canonical SMAD pathway and inducing translocation of the SMAD1/4/5/9 complex.^27^ SMAD7 inhibits both TGF-β and BMP-mediated SMAD signaling pathways. Here we show that Noregen inhibits BMP-2 and TGF-β activities as well as human norrin. We showed that Noregen decreases basal levels of SMAD signaling in a dose-dependent manner and inhibits TGF-β1-induced SMAD activity.

In summary, our recombinant norrin-derived growth factor, Noregen, retains the multi-functionality of human norrin: 1) endothelial barriergenesis; 2) retinal vascular regeneration; 3) neurogenic pathway activation; and 4) anti-fibrosis. Here we show that Noregen mimics the multiple activities of human norrin and is equivalent or superior in these functions. We believe this to be a potential novel treatment for ischemic retinopathies.

## Notes

### Competing Interest Statement

The authors have declared no competing interest.

### Funding

- NIH Common Fund: 2R44EY030807_02

